# Distinct *in vivo* dynamics of excitatory synapses onto cortical pyramidal neurons and inhibitory interneurons

**DOI:** 10.1101/2021.04.04.438418

**Authors:** Joshua B. Melander, Aran Nayebi, Bart C. Jongbloets, Dale A. Fortin, Maozhen Qin, Surya Ganguli, Tianyi Mao, Haining Zhong

**Author notes:** These authors contributed equally (Joshua B. Melander, Aran Nayebi). Correspondence should be addressed to S.G., T.M., or H.Z. Lead Contact: Haining Zhong, Ph.D., Vollum Institute, Oregon Health & Science University, 3181 SW Sam Jackson Park Road, L474, Portland, Oregon 97239, U.S.A., /.

## Abstract

Cortical function relies on the balanced activation of excitatory and inhibitory neurons. However, little is known about the organization and dynamics of shaft excitatory synapses onto cortical inhibitory interneurons, which cannot be easily identified morphologically. Here, we fluorescently visualize the excitatory postsynaptic marker PSD-95 at endogenous levels as a proxy for excitatory synapses onto layer 2/3 pyramidal neurons and parvalbumin-positive (PV+) inhibitory interneurons in the mouse barrel cortex. Longitudinal *in vivo* imaging reveals that, while synaptic weights in both neuronal types are log-normally distributed, synapses onto PV+ neurons are less heterogeneous and more stable. Markov-model analyses suggest that the synaptic weight distribution is set intrinsically by ongoing cell type-specific dynamics, and substantial changes are due to accumulated gradual changes. Synaptic weight dynamics are multiplicative, i.e., changes scale with weights, though PV+ synapses also exhibit an additive component. These results reveal that cell type-specific processes govern cortical synaptic strengths and dynamics.

## INTRODUCTION

The formation, plasticity, and rewiring of synaptic connections are fundamental to the function of neuronal circuits. Towards understanding these processes, the organization and dynamics of excitatory synapses onto cortical pyramidal neurons have been studied extensively using dendritic spines as the proxy (Bhatt et al., 2009; Grutzendler et al., 2002; Holtmaat and Svoboda, 2009; Trachtenberg et al., 2002). *In vivo* two-photon imaging of dendritic spines on fluorescent protein (FP)-expressing neurons has revealed several key principles governing these synapses: their strengths are log-normally distributed and change in a multiplicative manner (i.e., the magnitude of synaptic weight changes tends to be proportional to the size of the synapse) (Buzsáki and Mizuseki, 2014; Loewenstein et al., 2011; Ziv and Brenner, 2018). Furthermore, the addition and elimination of spiny synapses correlate with, and likely underlie, the acquisition of certain learned behaviors (Hayashi-Takagi et al., 2015; Hofer et al., 2009; Holtmaat et al., 2006; Johnson et al., 2016; Xu et al., 2009; Zuo et al., 2005a). These observations are foundational to our current experimental and theoretical understanding of brain function (Buzsáki and Mizuseki, 2014).

Although *in vivo* imaging of spine morphology has provided the first insights regarding synaptic dynamics *in vivo*, it is limited in two aspects. First, the use of spine morphology alone precludes the study of a major class of excitatory synapses – those residing directly on dendritic shafts (Anderson and Martin, 2006; Bozhilova-Pastirova and Ovtscharoff, 1996; Fiala and Harris, 1999). These synapses cannot be identified using neuronal morphology under light microscopy (Goldberg et al., 2003; Keck et al., 2011; Sancho and Bloodgood, 2018). Shaft excitatory synapses constitute the majority of excitatory synapses onto cortical inhibitory interneurons, such as parvalbumin-positive (PV+) interneurons (Harris and Shepherd, 2015; Huang, 2014; Kim et al., 2017; Lee et al., 2012). Understanding these synapses is critical, as normal brain function relies on an exquisite balance between excitation and inhibition (E/I) (Antoine et al., 2019; Xue et al., 2014; Zhou et al., 2014). Yet, to date, little is known about the distribution and dynamics of these synapses, particularly *in vivo*. Second, for synapses onto excitatory pyramidal neurons, a fraction of excitatory synapses do not reside on dendritic spines (Knott et al., 2006; Santuy et al., 2018). In addition, while spine size is correlated with synaptic size and strength, such correlations are not perfect (Fortin et al., 2014; Harris and Stevens, 1989; Matsuzaki et al., 2001 see also Figure 1G). Furthermore, spines that protrude axially from the parental dendrite are difficult to distinguish due to the poor axial-resolution of light-microscopy. Thus, the conclusions based on spine morphology alone can benefit from validation with methods that directly visualize synapses.

**Figure 1:**
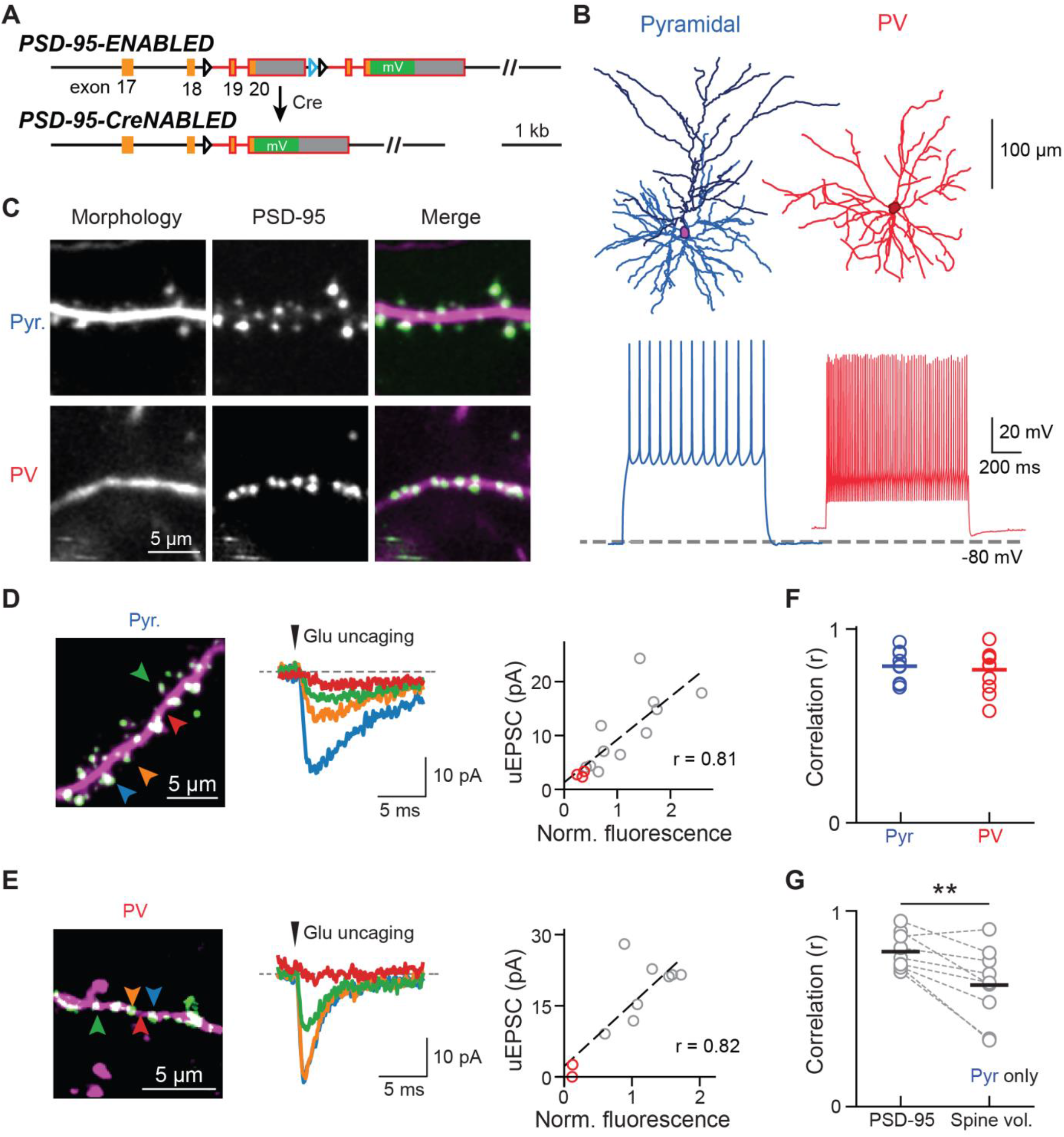
Visualizing endogenous PSD-95 as the proxy for the presence and weight of excitatory synapses onto Pyr and PV+ dendrites. (A) Schematic of the ENABLED/CreNABLED strategy for tagging endogenous PSD-95 with mVenus in a Cre-dependent manner. (B) Representative cellular reconstructions (top), and spike-trains (bottom) elicited via supra-threshold current steps from Pyr (left, blue) and PV+ (right, red) neurons. (C) Representative two-photon images of dendritic stretches from Pyr neurons (top) and PV+ inhibitory interneurons (bottom). PSD-95 (green) and dendritic morphology (magenta) were labeled simultaneously. (D, E) Two-photon glutamate uncaging experiment for Pyr (D) and PV+ (E) neurons in acute brain slices. Left: representative images showing a subset of uncaging locations (arrowheads) whose colors correspond to the traces shown. Middle: Example uEPSCs. Right: Correlation between integrated PSD-95^mVenus^ fluorescence intensities with uEPSC of the representative dendrite shown in the left panel. Red circles indicate stimulations at dendritic shaft locations without PSD-95^mVenus^ puncta. (F) Collective correlation (Pearson’s r) between PSD-95^mVenus^ fluorescence and uEPSC amplitude for Pyr and PV+ neurons. Each data point represents a different dendrite. Averages are: 0.81 ± 0.03, n (uncaging sites/dendrites) = 111/9 for pyramidal neurons; and 0.79 ± 0.03, n = 82/11. (G) The correlation between PSD-95^mVenus^ fluorescence and uEPSC amplitudes versus spine volume and uEPSC amplitudes in Pyr neurons. Only laterally-protruded spines were included. Averages are 0.79 ± 0.03 for PSD-95^mVenus^ and 0.62 ± 0.06 for spine volume, n (spines/dendrites) = 98/9. One-sided Wilcoxon signed-rank test: p = 0.004, W = 44. See also Figure S1.

To overcome these limitations, we used a molecular indicator to identify shaft excitatory synapses in interneurons and to visualize synaptic structure directly *in vivo*. An essential postsynaptic scaffolding protein at excitatory synapses, PSD-95 (Sheng and Kim, 2011), tagged with a FP, has been overexpressed to visualize synapses *in vivo* (Cane et al., 2014; Gray et al., 2006; Sun et al., 2019; Villa et al., 2016). However, the overexpression of PSD-95 has been shown to result in aberrant synaptic function, and defective synaptic plasticity (Béïque and Andrade, 2003; Ehrlich and Malinow, 2004; Elias et al., 2008; Schnell et al., 2002), making this approach less than ideal for studying synapses.

Herein, we used a conditional knock-in strategy called <u>en</u>dogenous l<u>ab</u>eling via <u>e</u>xon <u>d</u>uplication (ENABLED) to label PSD-95 at its endogenous levels with the yellow FP mVenus in neuronal subsets (Fortin et al., 2014). This allowed us to visualize excitatory synapses onto both layer 2/3 (L23) cortical spiny pyramidal (Pyr) neurons and aspiny PV+ inhibitory interneurons. The abundance of PSD-95 at a synapse correlated with the synapse’s sensitivity to the neurotransmitter glutamate more strongly than the spine volume did, suggesting that PSD-95 abundance provides a more accurate assessment of synaptic weight. Moreover, chronic *in vivo* two-photon imaging in the mouse barrel cortex revealed that the distribution and dynamics of excitatory synapses were cell type-specific. Although excitatory synapses onto both pyramidal neurons and PV+ interneurons both exhibited log-normal weight distributions, those onto PV+ dendrites were more uniform (i.e., exhibiting a narrower distribution) and, on average, contained lower-levels of PSD-95 than those onto pyramidal neurons. Excitatory synapses onto PV+ dendrites were also more stable over the experimental timescale (weeks) than their pyramidal counterparts. Furthermore, although weight changes of excitatory synapses onto both neuronal types were largely multiplicative (i.e., the magnitudes of changes scaled with synaptic weights), changes in PV+ neurons also exhibited a significant additive, weight-independent component. Notably, our results indicated that large, rapid changes in synaptic weights were rare, and substantial changes were primarily the product of incremental changes that accumulated over time. These results provide the first analysis of the organization and dynamics of shaft excitatory synapses onto inhibitory interneurons *in vivo*, and shed light onto cell type-specific mechanisms for maintaining the synaptic basis of cortical E/I balance.

## RESULTS

### Cell type-specific visualization of synapses via endogenous PSD-95 labeling

In order to visualize excitatory synapses onto different populations of cortical neurons, we employed the *PSD-95-ENABLED* mouse (Fortin et al., 2014), in which PSD-95 is tagged by mVenus and expressed at endogenous levels in a Cre-dependent manner (Figure 1A). Synaptic function and plasticity are normal both before and after Cre recombination in this mouse (Fortin et al., 2014). To simultaneously visualize neuronal morphology, we crossed *PSD-95-ENABLED* mice with *Ai9*, a Cre-dependent tdTomato reporter line (Madisen et al., 2010). Cre was introduced into L23 pyramidal neurons sparsely via *in utero* injection of adeno-associated virus (AAV) into *PSD-95-ENABLED;Ai9* double heterozygous mice, and into PV+ neurons by further crossing with the *PV-IRES-Cre* mouse (Jax #008069) (Hippenmeyer et al., 2005) to yield *PSD-95-ENABLED;Ai9;PV-IRES-Cre* triple heterozygous mice. Immunohistochemical experiments showed that *PV-IRES-Cre* mice reliably and specifically expressed Cre in PV+ neurons in the barrel cortex of adult mice (Figure S1A) (see also Pfeffer et al., 2013). For both strategies, labelled neurons exhibited morphological and electrical properties characteristic of each neuronal type (Figure 1B). PSD-95^mVenus^ expressed at endogenous levels can be imaged *in vivo* via a cranial window with high contrast in both neuronal types in the mouse barrel cortex (Figure 1C), with the cortical location verified using functional intrinsic-signal imaging (Figure S1B).

### PSD-95^mVenus^ puncta represent excitatory synapses

To validate that PSD-95^mVenus^ puncta represented functional excitatory synapses, we performed whole-cell patch-clamp recording of labeled neurons in acute brain slices while focally uncaging glutamate onto individual puncta in Pyr and PV+ dendrites using a two-photon microscope. Uncaging immediately adjacent to PSD-95^mVenus^ puncta triggered robust excitatory post-synaptic currents (uEPSCs) in both neuronal types (Figures 1D and 1E), which were blocked by the AMPA receptor antagonist NBQX (Figure S1C). In contrast, uncaging at dendritic locations in which PSD-95^mVenus^ puncta were absent gave little to no current (Figures 1D, 1E and S1D). Importantly, the uEPSC amplitude correlated strongly with the fluorescence intensity of the corresponding PSD-95^mVenus^ punctum for both neuronal types (Figures 1D, 1E, and 1F), indicating that PSD-95^mVenus^ intensity at a given punctum is an accurate measurement of the “weight” of the synapse, defined as the response to uncaged glutamate recorded at the soma.

In pyramidal neurons, synaptic weight has been shown to correlate with spine volume (Matsuzaki et al., 2001), and our data also indicated that PSD-95^mVenus^ intensity correlated with the volume of clear, laterally-protruding spines (Figure S1E). However, many synapses overlapped with the dendritic shaft (e.g., Figure 1C), making measurements of the spine volume difficult. Additionally, the orientation of spines may change over time, which can result in errors when tracking the same synapses chronically using morphology alone (e.g., Figure S1F). Furthermore, for clear laterally-protruding spines, uEPSCs were better correlated with integrated PSD-95^mVenus^ fluorescence than spine volume, suggesting that the abundance of PSD-95 was a more reliable measure of synaptic weight (Figure 1G). Taken together, these data indicate that PSD-95^mVenus^ puncta label functional glutamatergic synapses onto both L23 pyramidal neurons and in PV+ interneurons, and that PSD-95^mVenus^ fluorescence intensity can be used to estimate a synapse’s weight more accurately than spine volume can.

### The weight distribution of excitatory synapses is cell type-specific

We asked whether the distribution of excitatory synaptic weights onto the two cell types, as inferred from PSD-95^mVenus^ intensity, differed from each other *in vivo*. *S*egments of Pyr and PV+ dendrites residing in layer 1 (depth <100 µm from pia) of the barrel cortex (S1) of adult mice were imaged (Figure 1C). Some PV+ dendrites lacked PSD-95^mVenus^ expression in their entirety, presumably due to incomplete Cre-recombination efficiency in the *PSD-95-ENABLED* mice (Fortin et al., 2014), and were not further examined. Imaged dendritic stretches were then manually scored to identify and classify each PSD-95^mVenus^ puncta.

Dendrites from L23 pyramidal neurons exhibited a higher synapse density than those of PV+ dendrites (Pyr: 1.02 ± 0.03 synapse/µm, PV+: 0.87 ± 0.01 synapse/µm; two-sided t-test: *t* = 2.9, p < 0.01) (Figure 2A). In L23 pyramidal neurons, PSD-95^mVenus^ was nearly exclusively present as punctate signals, and most (∼96%) spines contained PSD-95^mVenus^ puncta. Importantly, PSD-95^mVenus^ allowed for the visualization of a large number of synapses that colocalized with the dendritic shaft in Pyr dendrites (∼1/3 of all puncta). These shaft-colocalized synapses would have been omitted when imaging morphology only. They likely include both axially-protruding spines and shaft excitatory synapses, which make up about 20% of total excitatory synapses in the cortex, as reported in electron microscopy (EM) studies (Fiala and Harris, 1999; Knott et al., 2006; Santuy et al., 2018). In contrast, excitatory synapses onto PV+ dendrites were mostly restricted to the dendritic shaft (Figures 1C and 2A), consistent with previous studies (Goldberg et al., 2003; Keck et al., 2011; Sancho and Bloodgood, 2018). Nevertheless, small spines enriched with PSD-95^mVenus^ can be occasionally found on otherwise aspiny PV+ dendrites (Figure S1G) (see also Keck et al., 2011; Sancho and Bloodgood, 2018).

**Figure 2:**
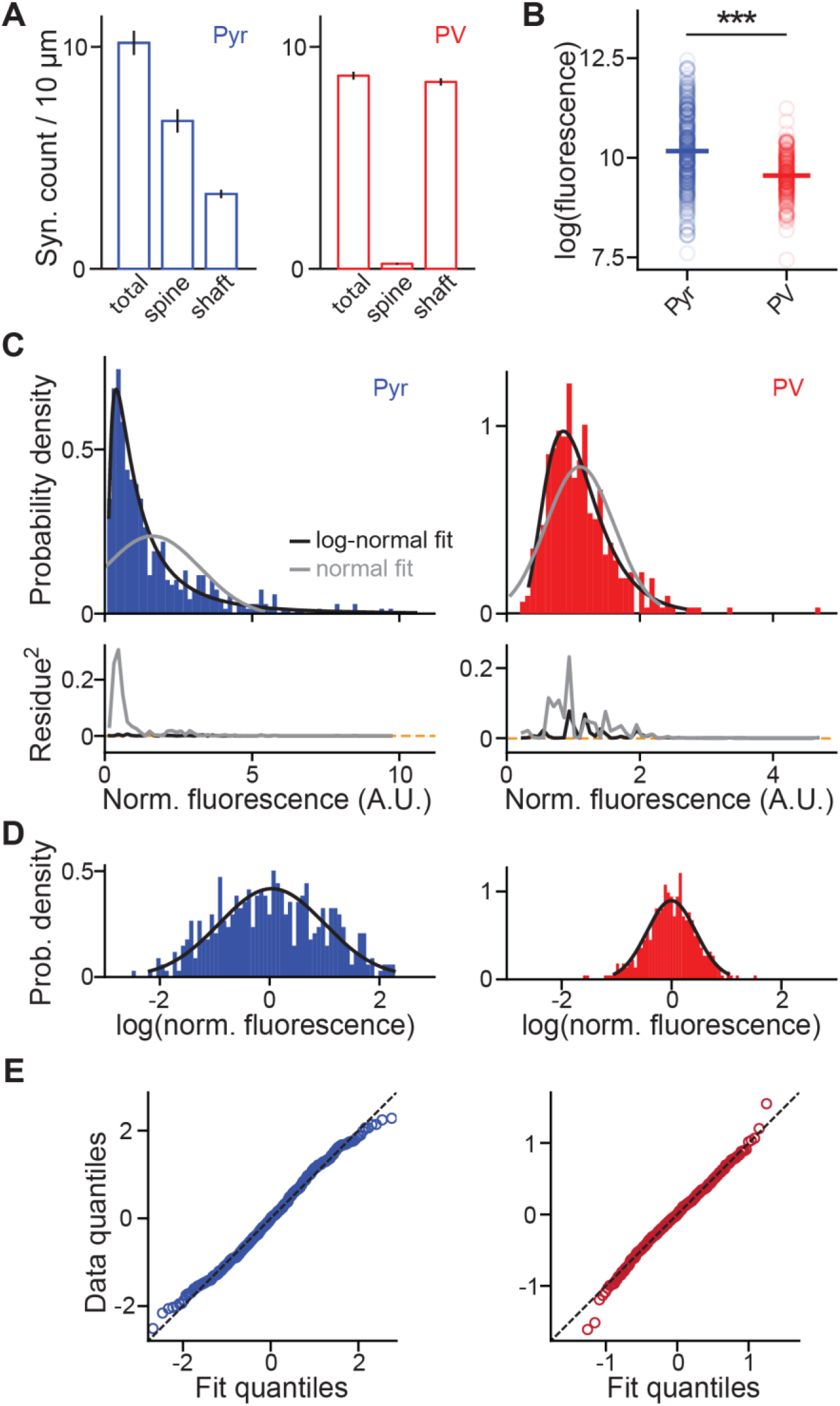
Population characteristics of the organization of synapse and their weights. (A) Density of PSD-95^mVenus^ puncta, and the fractions of laterally-protruding spines and of those colocalized with dendritic shafts, on Pyr (left, blue) and PV+ (right, red) dendrites in layer 1 *in vivo*. (B) Absolute integrated PSD-95^mVenus^ fluorescence of synapses on pyramidal and PV+ dendrites in the logarithmic scale. Only synapses between 10 – 50 µm beneath the pial surface are selected for this analysis to minimized depth dependent variations in imaging conditions. Medians are 10.3 ± 0.06 (s.e.m.), n = 303 (synapses) for pyramidal, and 9.5 ± 0.04, n = 191 for PV. Wilcoxon rank-sum test: p < 0.001, U = 8.28. (C) Probability density histograms of normalized synaptic PSD-95^mVenus^ fluorescence fitted by log-normal or normal distributions (top) and their squared residues (bottom) for pyramidal and PV+ neurons, as indicated. n = 452 and 409 for pyramidal and PV+, respectively. (D, E) Histograms of the logarithm of synaptic PSD-95^mVenus^ fluorescence plotted with the best-fit normal distribution (D) and their quantile-quantile plots (E). Note that the distribution of weights is broader for pyramidal (left, blue) than for PV + (right, red) dendrites.

We next compared the weights of synapses onto both neuronal types by quantifying the integrated fluorescence of individual PSD-95^mVenus^ puncta. Synapses onto L23 pyramidal neurons were significantly brighter than those onto PV+ neurons, suggesting that excitatory synapses onto pyramidal neurons are more enriched for PSD-95 than those onto PV+ interneurons (Figure 2B). It is important to note that this result does not necessarily mean that PV+ synapses are weaker than pyramidal synapses because, while our results indicate that the PSD-95^mVenus^ can be used to infer weight within a cell type (Figures 1D and 1E), the relationship between intensity and synaptic strength may differ across cell types. For both cell types, the distribution of synaptic weights was best fit by a log-normal distribution, but not a normal distribution (Figures 2C – 2E). At the same time, the intensity distribution of pyramidal synapses was much broader (full width half maximum of fit in log scale ± 95% confidence level: Pyr, 2.5 ± 0.8; PV+, 0.97 ± 0.08; Figure 2D), suggesting a wider range of synaptic weights onto L23 pyramidal neurons than onto PV+ interneurons. These results indicate that the composition and organization of excitatory synapses are cell type-dependent.

### Excitatory synapses onto PV+ interneurons are stable at the month time-scale

Previous studies imaging dendritic spines suggested that a fraction of cortical spiny synapses are added or eliminated over behaviorally-relevant timescales (Holtmaat et al., 2006; Xu et al., 2009; Zuo et al., 2005a). More recently, aberrant synaptic dynamics were found to be associated with various neurological disorders and neurodegenerative diseases (Guo et al., 2015; He et al., 2017; Murmu et al., 2015; Spires et al., 2005; Tsai et al., 2004). We asked whether the dynamics of excitatory synapses differed between the two neuronal types by longitudinally imaging both L23 pyramidal and PV+ dendrites every four days for 7 – 8 time points in the barrel cortex (Figures 3A and 3B). To track the dynamics of a given synapse precisely, we developed custom analysis software in MATLAB that permitted the side-by-side comparison and annotation of the same stretch of dendrite over all time points simultaneously (Supplemental Figure 2). Each synapse was manually scored for its persistence, addition, or elimination across all time points (e.g., Figure 3B; see **STAR METHODS** for details).

**Figure 3.**
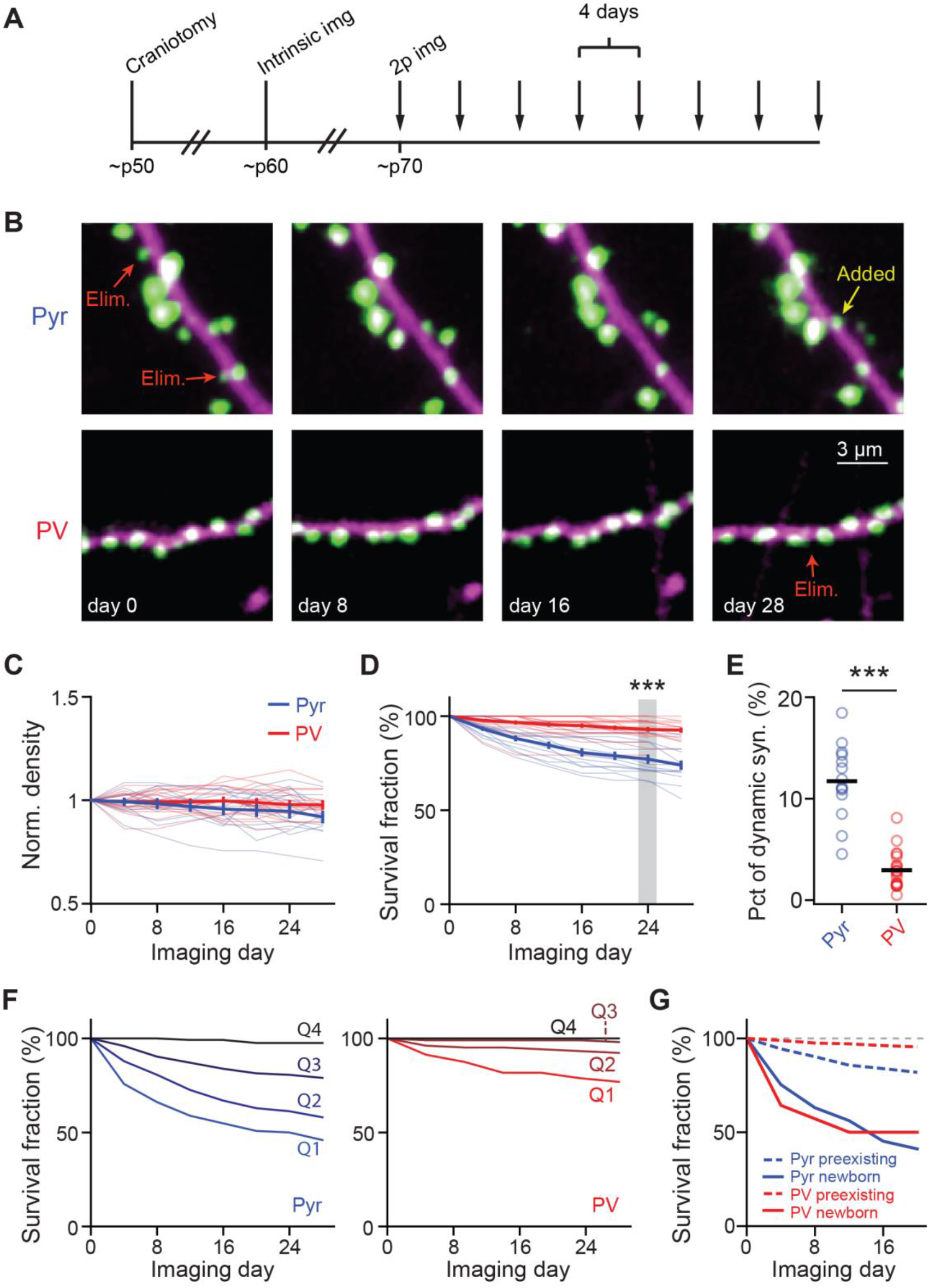
Synapses onto PV+ dendrites are more stable compared to those onto pyramidal dendrites. (A) Experimental protocol for longitudinal in-vivo experiments. (B) Representative images of dendritic stretches of Pyr (top) and PV+ (bottom) neurons in layer 1 imaged longitudinally for a month. Every other time point is shown. Arrows indicate representative added, eliminated (Elim.), or persistent synapses. (C) Time course of the synaptic density of Pyr and PV+ dendrites normalized to day 0. (D) Survival fraction of synapses present on the first imaging session at subsequent time points. At 24 days, only 77 ± 2% remain on pyramidal dendrites, compared to PV+ 93 ± 1% on PV+ dendrites. n (synapses/dendrites/animals) = 594/14/4 for pyramidal, and 638/21/7 for PV. Wilcoxon rank-sum test: p < 0.001, U = 2.09. (E) Average percentage of all synapses on a dendritic stretch that were added or eliminated during a 4-day interval. Averages are (Pyr: 11 ± 1%, PV: 3 ± 0.4%; Wilcoxon rank-sum test: p<0.001, U=4.8). (F) Survival fractions of four quartiles (from weak to strong: Q1 – Q4) of synapses sorted by weight on the first imaging day for pyramidal (left) and PV+ (right) dendrites. (G) Survival fractions of newborn versus preexisting synapses onto pyramidal (blue) and PV+ (red) dendrites as indicated. n = 552, 441, 73 and 14, respectively, for Pyr preexisting, PV preexisting, Pyr newborn, and PV newborn. See also Figures S2 and S3.

Synapses on the two neuronal types exhibited distinct patterns of structural plasticity. As expected, the total number of synapses observed on each dendritic segment was relatively stable throughout the month of imaging, despite ongoing synaptic additions and eliminations (Figure 3C). Consistent with previous studies (Holtmaat et al., 2005; Ma et al., 2016; Sun et al., 2019; Tjia et al., 2017), the dendrites of L23 pyramidal neurons exhibited a moderate level of synaptic addition and elimination (Figures 3D and 3E). Although most synapses were stable, 23 ± 2% of synapses on Pyr dendrites observed at day 0 were lost at day 24 (Figure 3D). Between each four-day imaging window, 11 ± 1% synapses on average were added to or eliminated from Pyr dendrites (Figure 3E). PV+ dendrites, however, exhibited much lower levels of synaptic turnover (Figures 3D and 3E). At day 24, PV+ dendrites lost only 7 ± 1% of the synapses observed at day 0 (p < 0.001, cf. Pyr dendrites; Figure 3D). The percentage of dynamic synapses onto PV+ dendrites per four-day window was also more than 3-fold lower than that of Pyr dendrites (Figure 3E).

Previous results have suggested that larger spines tend to be more stable, and that newborn spines are more likely to be eliminated than preexisting spines (Holtmaat et al., 2005; Majewska et al., 2006; Xu et al., 2009; Zuo et al., 2005b). We first asked whether synapses with higher PSD-95^mVenus^ content were more stable. Synapses from each neuronal type were grouped into four quartiles based on their integrated PSD-95^mVenus^ fluorescence on the first imaging day. As assayed using the survival function, smaller synapses were more dynamic than larger ones in both neuronal types (Figure 3F). For all quartiles, PV+ synapses were more stable than pyramidal synapses (Figure 3F). Even for pyramidal dendrites, the synapses in the upper-most quartile were stable, exhibiting little turnover over the course of the experiment. In addition, we found that newborn synapses in both neuronal types were much more likely to be eliminated than preexisting synapses on the same dendrites (Figure 3G). Altogether, these results indicate that the dynamics of synaptic addition and elimination are cell type-specific *in vivo*, with shaft excitatory synapses onto interneurons being much more stable than those onto Pyr neurons. Furthermore, the synaptic turnover events in both neuronal types are dominated by smaller and new-born synapses, with stronger and persistent synapses exhibiting a high degree of stability.

### Synaptic weight dynamics are cell type-specific

We next move beyond observing the binary addition and elimination of synapses, to observing the more detailed evolution of the weights of individual synapses over time, which has been less studied *in vivo*. In order to do so, we pooled data from different dendrites and days together by normalizing all synapses along a dendrite at each time point to the average intensity within the 40^th^ through 60^th^ percentiles of those synapses. This normalization was necessary to correct for inevitable variations in imaging conditions across dendrites, animals, and days (e.g., see Figure S3A and S3B), and appeared more robust than an alternative approach that normalized green fluorescence intensity to the cytosolic reporter tdTomato due to the differential bleaching of PSD-95^mVenus^ and tdTomato signals (Figure S3C and S3D).

We asked whether synaptic weight changes were cell type-specific. Only synapses from the 25^th^ to 75^th^ percentiles were analyzed to minimize the effect of relatively low signal-to-noise ratios associated with the smallest synapses. For both neuronal types, the average weight of synapses that persisted throughout all imaging time points did not change over time (Figure 4A). This lack of change was presumably due to synaptic weight increases and decreases across synapses canceling each other out. Indeed, when the absolute weight changes were analyzed (Figure 4B), the weights of individual synapses deviated from their original weight substantially. The degree of weight change, however, was cell type specific. Weight changes of PV+ synapses were moderate (0.24 ± 0.02 units on the natural log scale at day 24, corresponding to ∼27% changes; Figure 4A). In contrast, persistent Pyr synapses exhibited greater changes (0.46 ± 0.03 units at day 24, corresponding to ∼58% changes; Figure 4A).

**Figure 4:**
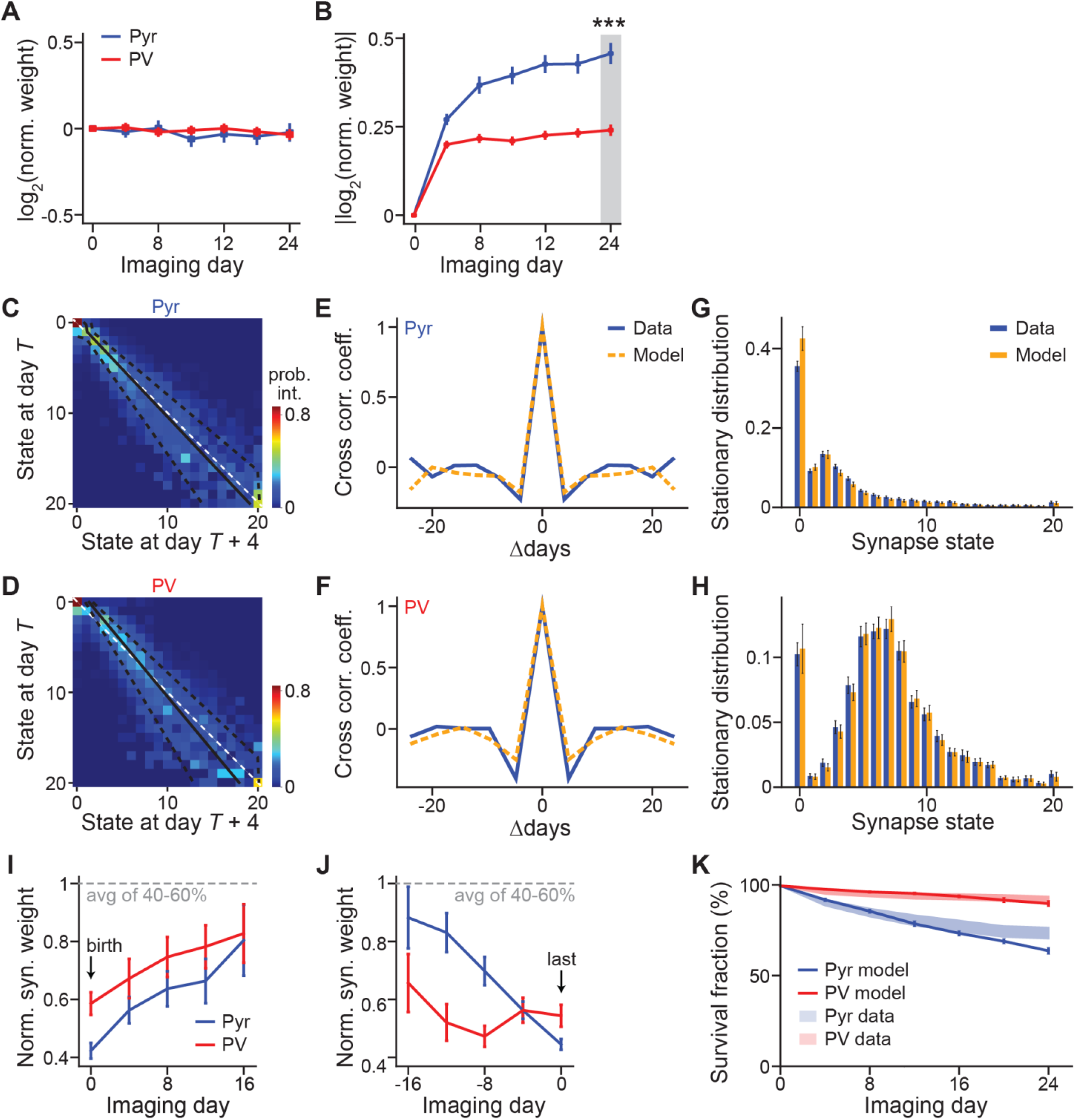
Markov-chain model reveal cell type-specific principles of synaptic weight dynamics. (A, B) Average of signed (A) and absolute (B) synaptic weight changes relative to day 1 in the natural log scale. The log scale was used so that increase and decrease in synaptic weights are equal in the analysis. n = 174 for Pyr and 190 for PV+. (C, D) The Markov state transition matrices of synapses onto both pyramidal (C) and PV+ (D) dendrites exhibit a central band about the unity line (white dashed) that widens as current synaptic strength increases. Each row represents the conditional future weight distribution at the next time point (x axis) given a current synaptic weight bin (y axis). Synaptic strength increases with bin index. For comparison, the mean of each row, or conditional future weight distribution (solid black line) and its ± 1 standard deviation (dashed black lines) as determined by a fitted Kesten process (see Methods) are superimposed on the Markov transition matrix. (E, F) Cross correlation coefficient of synaptic weight changes. Graphs denote the correlation coefficient between weight changes from day T to day T+4 and weight changes from day T+Δ to day T+Δ+4, averaged over all synapses and days T (see Methods). Dashed orange curves indicate weight change correlations for our models’ Markovian dynamics which, as expected, are close to be 0 as soon as Δ is nonzero. Blue curves indicate weight change correlations in the data are similarly weak at nonzero Δ indicating the validity of our Markovian modelling, at least on a time scale where observations are taken every 4 days. (G, H) Steady state distributions of synaptic strength for both synapse classes as computed from experimental data (orange) and as predicted by our model (blue). Error bars denote standard deviation across 30 bootstrap runs (see Methods). (I, J) Addition-triggered (I) and elimination-triggered (J) averages of weight-trajectories for both synapse classes types, aligned to birth and death, respectively. Error bars denote standard error of the mean. (K) The survival fraction as a function of time predicted by the Markovian transition model (solid lines) and the actual survival fraction computed in the data (shades) for both synapses onto pyramidal (blue) and PV+ (red) neurons. Error bars and the width of the shaded curves both indicate standard error of the mean.

### Markov model of weight-dynamics predicts cell type-specific stationary synaptic weight distributions

We then modeled the dynamics of excitatory synapses onto both neuronal types as Markovian processes. The synaptic weights were binned into 20 states, with one further state added to represent strength zero (i.e., not present), yielding a 21X21 Markov state transition matrix (Figures 4C and 4D) that described the synaptic weight change between adjacent time points (see **Star Methods**). Each row of the matrix denoted the probability distribution of synaptic weights on the next observation day for synapses within a binned weight range on the current observation day. To test the validity of our Markov assumption, we examined whether changes in synaptic weights from day T to day T+4 were correlated with changes in synaptic weights from day T+Δ to day T+Δ+4 (imaging was performed once every 4 days). The cross-correlation of synaptic weight changes between two pairs of observations days dropped sharply as the separation Δ between those pairs of observation days increased for both pyramidal and PV+ dendrites (Figures 4E and 4F) both in the data and in our model. In other words, knowledge of synaptic weights 8 days ago did not provide more information about the current weights, as measured via linear correlation, than did knowledge of the weights 4 days ago. These results indicate that the underlying dynamics of synaptic strengths can be modelled as a Markovian process on the time scale of our observations.

Above, we found that synaptic weights adopted a cell type-specific log-normal distribution (Figures 2C and 2D). However, the underlying mechanism was poorly understood. Our Markov model of synaptic dynamics predicts a particular steady-state distribution of synaptic weights, which is equivalent to iterating the dynamics over time from any initial distribution, mimicking the process of synaptic evolution. We computed this predicted stationary distribution and found that it was very similar to the empirically measured distribution for both neuronal types (Figures 4G and 4H). Thus, the Markovian state transition matrix, while fit using only pairs of synaptic weights from adjacent time points, can be generalized to accurately predict the steady-state distribution of synaptic weights, suggesting that the synaptic weight distribution is intrinsically determined by the dynamics of synapses onto each neuronal type.

### Gradual accumulative changes predict synapse addition and elimination

A noticeable feature of the Markov state transition matrices was that synaptic changes were gradual, with very few large jumps in synaptic weights, as evidenced by the concentration of higher transition probabilities near the diagonal in Figures 4C and 4D. One prediction of such Markovian dynamics is that synapses are both born into and die in states of weak synaptic strengths. To test this prediction, we examined the dynamics of synaptic weights aligned by their addition or elimination day. For both cell types, new synapses were added with low synaptic weights, with those that survived gradually gaining strength over weeks (Figure 4I). At the same time, synapses were eliminated from a low synaptic weight state (Figure 4J). These results together suggest that cortical synaptic changes, if they happen, are gradual, and large changes (e.g., the formation or loss of a strong synapse) arise primarily through the gradual accumulation of multiple small events unfolding over the course of weeks.

To further test the ability of our Markov model to make long range predictions about synapse elimination, we examined whether it could predict the fraction of synapses that survived out to 28 days. We did so by iterating our model over multiple observation periods and found that it matched well the empirical survival function for synapses onto both neuronal types (Figure 4K). Moreover, both the model and the empirical data revealed a much higher synaptic turnover rate (lower survival fraction) in L23 pyramidal dendrites than in PV+ dendrites (Figure 4K). Overall, these results indicate that our Markovian synaptic dynamics model accurately predicts a fundamental cell type-specific difference that emerges over a month-long time scale.

### Synaptic weight changes are not solely multiplicative

Our data provides the opportunity to test two opposing models of synaptic weight dynamics, additive versus multiplicative dynamics (Loewenstein et al., 2011). In the additive model, unobserved pre- and post-synaptic activity leads to a synaptic change that is independent of the current weight of the synapse. In contrast, the multiplicative model hypothesizes that the same unobserved activity leads to a change whose magnitude is proportional to the current weight. These models can thus be differentiated by examining how changes in synaptic weight scale with the current weight.

The Markov state transition matrix of both pyramidal and PV+ dendrites exhibited an increasingly-widened central band as the previous day’s synaptic weights increased (Figures 4C and 4D). This widening band indicates that the average magnitude of synaptic change grows with existing synaptic strength, suggesting that the evolution of synaptic weights in both neuronal types has a multiplicative component. This prediction is further supported by fitting the synaptic weights with a more constrained Kesten process consisting of both additive and multiplicative components (Hazan and Ziv, 2020; Statman et al., 2014; Ziv and Brenner, 2018). Under such a framework, the intercept of the linear-fit to weight changes (w_t+1_ – w_t_) would indicate an additive component, whereas a linear correlation between the variance of synaptic weight changes with the square of the current synaptic weight indicates a multiplicative component (Figure S4). The fit Kesten process produced a widening central band structure with increasing state, which can also be seen from the Markov transition matrix distributions (overlaid black lines in Figures 4C and 4D), indicating consistency between the more constrained Kesten model and the more general Markov transition model. Synapses onto both cell types have a clear multiplicative component (Figure S4B). At the same time, as evidenced by deviation from the diagonal line (Figure 4D) and the positive y-intercept (Figure S4A), the Kesten model fit to synapses onto PV+ cells also revealed a significant additive component. Such an additive component was less evident for pyramidal synapses (Figure 4C and S4A). Indeed, removing the additive component from the Kesten model resulted in a poor fit for PV+ synapses (r^2^ = 0.03 and 0.89 for without and with the additive component, respectively), but only moderately affected the fit for pyramidal synapses (r^2^ = 0.53 and 0.62 for without and with the additive component, respectively). Together, these data suggest that, although excitatory synaptic weight changes are multiplicative, synapses onto PV+ dendrites also have a significant additive component.

## DISCUSSION

Here, we present, to our knowledge, the first longitudinal *in vivo* imaging of a major postsynaptic structural protein, PSD-95, in the barrel cortex, without protein overexpression. Using PSD-95^mVenus^ puncta as the marker for endogenous excitatory synapses, this study also presents the first *in vivo* description of the dynamics of a major cortical synapse type, excitatory shaft synapses onto inhibitory interneurons. We found that synaptic organization and plasticity are cell type-specific. Furthermore, by using PSD-95^mVenus^ fluorescence intensity as an estimate for the weight of a synapse, together with a quantitative Markov transition model, we illustrated several principles of synaptic dynamics that may have implications for both the experimental and computational understanding of synaptic plasticity. First, the stationary synaptic weight distribution is cell type-specific and is intrinsically associated with the day-to-day synaptic dynamics. Second, the majority of synaptic weight changes in the cortex is incremental. Third, although synaptic weight changes follow multiplicative dynamics in both cell types, there is a cell type-specific additive component present only in PV+ dendrites.

Synaptic marker proteins of both excitatory and inhibitory synapses have been previously imaged *in vivo* (Cane et al., 2014; Chen et al., 2012; Gray et al., 2006; van Versendaal et al., 2012; Villa et al., 2016). However, these studies relied on overexpressed synaptic proteins tagged by an FP. Overexpression causes undesirable effects, making these results difficult to interpret at times. Specifically, the overexpression of PSD-95 is known to alter synaptic function and occlude long-term potentiation. A major gap therefore remains in our understanding of the organization and dynamics of endogenous synaptic proteins *in vivo*. The ENABLED strategy provides a viable way to visualize endogenous proteins that can potentially be expanded to other synaptic proteins. Currently, there are also parallel efforts to develop alternative strategies to label endogenous proteins for live imaging (Mikuni et al., 2016; Nishiyama et al., 2017; Suzuki et al., 2016; Zhong et al., 2020). Endogenous protein imaging will likely constitute an important research avenue in the future. At the same time, endogenous proteins are usually expressed at much lower levels than overexpressed proteins, bringing additional challenges that call for maximizing the signal and reducing the noise, such as FP improvements and microscopy optimization.

Previous studies of synaptic dynamics *in vivo* using spine imaging have generated important insights. One limitation of using spine morphology is the inherent difficulty in seeing axially protruding spines and shaft excitatory synapses that also exist in cortical pyramidal neurons (Fiala and Harris, 1999; Knott et al., 2006; Santuy et al., 2018). This is largely overcome in our study by visualizing the synaptic molecular contents together with morphology. Our results confirm several previous conclusions from spine imaging, including the log-normal distribution and multiplicative dynamics of spiny synapses. A log-normal distribution of synaptic weights suggests that a small percentage of larger synapses may provide a critical ensemble of neuronal connections, while the smaller synapses provide fine adjustment to the circuit and repertoire for additional plasticity and recruitment of new circuits (Buzsáki and Mizuseki, 2014). Interestingly, the largest synapses in both neuronal types were rarely eliminated (Figure 3F), possibly reflecting that the critical neuronal network formed by strong synapses is highly conserved over time. Additionally, although previous studies on *in vivo* synapse dynamics were mostly focused on the binary addition and elimination of spines, we demonstrate that there is rich information within the synaptic weight dynamics. Our results also suggest that weight dynamics are well described by small, analog changes, and larger weight changes and synaptic additions and eliminations are the cumulative result of these smaller events. These results call attention to such analog changes that should be explored in future studies.

Compared to previous studies, PSD-95 labeling allows us to examine *in vivo* an understudied, major class of excitatory synapse – the shaft excitatory synapses onto inhibitory interneurons. Interestingly, these synapses exhibit markedly different characteristics compared to the spiny synapses onto L23 pyramidal neurons. Shaft excitatory synapses onto PV+ neurons are packed at a lower density, contains a lower PSD-95 content, and exhibit a narrower range of synaptic weights. They are also less dynamic than the spiny synapses on pyramidal neurons. Although the exact mechanism underlying these differences is unclear, these observations add to the notion that plasticity in the cortex is highly specific and governed, in part, by the identity of the post-synaptic cell. The uniformity and stability of synapses onto PV+ neurons are consistent with the canonical function of interneurons as maintainers of stable E/I balance in the brain (Antoine et al., 2019; Xue et al., 2014; Zhou et al., 2014). In contrast, the larger dynamic range in synaptic weights and higher turnover of synapses onto pyramidal neurons may allow them to undergo plastic changes to rewire cortical circuits when needed, such as during behavioral adaptation, learning, and memory. In addition, the differences between spine and shaft synapses may be intrinsic to their geometric constraints. For example, spine protrusions effectively increase the cylindrical space along the dendrite, allowing more synapses to be formed per unit dendritic length. Similarly, spiny synapses enable the dendrite to sample a larger space and possibly interact with more potential presynaptic partners. Interestingly, the synaptic distribution is determined by the day-to-day dynamics of each synapse type. This may be the result of underlying homeostatic mechanisms (Turrigiano, 2012) that control the overall excitability of synapses onto both cell types and contribute to the cortical E/I balance.

Finally, classic models of synaptic plasticity often treat potentiation and depression events additively, namely through the addition of a fixed quantity that is independent of a synapse’s current strength (Gerstner et al., 1996; Hopfield, 1982; Song et al., 2000). However, recent studies (Loewenstein et al. 2011) suggest that *in vivo* spine size changes onto L23 pyramidal neurons might obey multiplicative dynamics, i.e., synaptic changes being scaled with the synapses’ current strengths. Here, by directly measuring the molecular content of the postsynaptic density and using it to assess synaptic weight, we find that that both Pyr and PV+ neurons exhibit multiplicative dynamics, suggesting that multiplicative scaling may be a general rule of synaptic plasticity. Interestingly, PV+ synapses, but not Pyr synapses, also exhibited an additive component, providing *in vivo* evidence that additive and multiplicative mechanisms are not mutually exclusive (Ziv and Brenner, 2018) and the degree of their co-existence is cell type-specific.

## SUPPLEMENTAL INFORMATION

Supplemental Information includes four figures.

## AUTHOR CONTRIBUTIONS

Conceptualization, J.B.M., T.M., & H.Z.; Methodology, J.B.M., and H.Z.; Software, J.B.M., A.N., and H.Z.; Investigation, J.B.M., A.N., B.C.J., D.A.F.; Formal Analysis, J.B.M., A.N., B.C.J., and H.N.; Resource, M.Q.; Writing – Original Draft, J.B.M., A.N., S.G., T.M., and H.Z.; Writing – Reviews & Editing, J.B.M., A.N., B.C.J., S.G., T.M., and H.Z.; Funding Acquisition, S.G., T.M., and H.Z.; Supervision, S.G., T.M., and H.Z.

## ACKNOWLEDGEMENTS

We thank Drs. Wenzhi Sun, Na Ji, and Vijay Iyer, and Mr. Daniel Flickinger at Howard Hughes Medical Institute Janelia Research Campus, Drs. Joseph Wekselblatt and Cris Niell at University of Oregon, Ms. Valerie Osterberg and Dr. Vivek Unni at Oregon Health & Science University, for help with surgeries, hardware, and software for *in vivo* two-photon imaging; Dr. Daniel O’Connor at Johns Hopkins University for intrinsic imaging setup; Dr. Emmeke Aarts at University of Utrecht for data analyses not included in this manuscript. We thank Drs. Lei Ma and Vivek Unni at Oregon Health & Science University and Yi Zuo at University of California Santa Cruz for critical comments for the manuscript. This work was supported by three NIH BRAIN Initiative awards (H.Z. and T.M.): U01NS094247, R01NS104944, and RF1MH120119, an NINDS R01 grant R01NS081071 (T.M.), an NINDS R21 grant R21NS097856 (H.Z.), awards from the Simons and James S. McDonell Foundations (S.G.), and an NSF CAREER award (S.G.).

## STAR METHODS

### RESOURCE AVAILABILITY

#### Lead Contact

Further information and requests for resources and reagents should be directed to and will be fulfilled by the Lead Contact, Haining Zhong.

#### Materials Availability

This study did not generate new unique reagents.

#### Data and Code Availability

The code generated for data analysis during this study will be made available at Github upon acceptance of the manuscript. The imaging data will be made available upon acceptance of the manuscript.

### EXPERIMENTAL MODEL AND SUBJECT DETAILS

Animal handling and experimental protocols were performed in accordance with the recommendations in the Guide for the Care and Use of Laboratory Animals of the National Institutes of Health and were approved by the Institutional Animal Care and Use Committee (IACUC) of the Oregon Health & Science University (#IS00002792).

#### Animals

Mice strains used in this study included: *PSD-95-ENABLED* (JAX #026092), *parvalbumin-IRES-Cre* (*PV-IRES-Cre*; JAX #008069), *Ai9* (JAX #007909). All mice were back-crossed to C57BL/6J (Charles River) background for at least five generations. *PSD-95-ENABLED;Ai9;PV-IRES-Cre* triple heterozygous mice were bred for simultaneously visualizing PSD-95 and cell morphology in PV+ interneurons. *PSD-95-ENABLED;Ai9* double heterozygous embryos were undergone in utero injection of AAV to their lateral ventricles to broadly and sparsely label a subset of neurons throughout the brain, including L23 pyramidal neurons in the barrel cortex. All mice were housed in standard laboratory conditions in groups of 2 or more with *ad libitum* access to food and water at all times inside a vivarium with a 12-hour light/12-hour dark cycle. Experiments were performed during both the light and dark cycles. For longitudinal studies of synapse dynamics *in vivo*, a glass window was installed to the mouse skull at p45-p65 via craniotomy. *In vivo* two-photon imaging was performed at least two weeks post window installation in both male and female mice under isoflurane anesthesia.

#### Acute Slice

Preparation of acute slices for two-photon glutamate uncaging and whole-cell patch-clamp recording was performed as previously described (Fortin et al., 2014). Briefly, coronal brain slices (300 µm thick) were prepared from adult mice typically at p21-p28 using ice-cold cutting solution containing the following (in mM): 110 choline chloride, 25 NaHCO_3_, 25 glucose, 2.5 KCl, 7 MgCl_2_, 0.5 CaCl_2_, 1.25 NaH_2_PO_4_, 11.5 sodium ascorbate, and 3 sodium pyruvate and being gassed with 95% O_2_/5% CO_2_. For experiments involving the *PV-IRES-Cre* driver line P56 – P59 mice were used since parvalbumin (PV) expression turns on relatively late during development (Bergmann et al., 1991; Nitsch et al., 1990). Slices were incubated in a recovery chamber with gassed artificial cerebrospinal fluid(ACSF) containing (in mM): 127 NaCl, 25 NaHCO_3_, 25 D-glucose, 2.5 KCl, 1 MgCl_2_, 2 CaCl_2_, and 1.25 NaH_2_PO_4_ at 34-36 °C for 30 minutes and then at room temperature (RT) for up to 8 hours. Individual slices were subsequently transferred to a recording chamber mounted on an upright microscope (Olympus BX51WIF) that was circularly perfused with freshly oxygenated ACSF at RT.

### METHOD DETAILS

#### Acute Slice Electrophysiology and Glutamate Uncaging

Whole-cell voltage-clamp recordings of uncaged excitatory post-synaptic currents (uEPSCs) from layer 2/3 neurons were recorded using an Axonpatch 200B amplifier (Molecular Devices) at RT. Electrophysiological signals were filtered at 2 kHz, and digitized and acquired at 20 kHz with custom software written in MATLAB. Borosilicate pipettes (2.8 – 6 MΩ; Warner Instruments) were filled with potassium gluconate-based internal solution containing (in mM): 120 potassium gluconate, 2 MgCl_2_, 0.1 CaCl_2_, 5 NaCl, 4 Na_2_-ATP, 0.3 Na-GTP, 15 Na-phosphocreatine. 10 HEPES, 1.1 EGTA, 0.01 Alexa-594 (Invitrogen), 3 mg/ml biocytin; pH 7.3; 290 mOsm. AMPA receptor mediated uEPSCs were isolated by addition of (in µM) 1 TTX, 10 SR 95531, 5 CPP in the ACSF to block voltage-gated Na channels, GABA-A receptors, and NMDA receptors, respectively. Cells were held at −70 mV throughout the course of the experiment.

*In vitro* two-photon imaging and uncaging was performed on a custom-built two-photon microscope controlled by ScanImage (Pologruto et al., 2003). Two Ti:sapphire lasers (MaiTai, Spectra Physics) were combined with polarized optics and passed through the same set of scan mirrors and objective for simultaneous imaging and uncaging. A water-immersion objective from Olympus (60X, 1.0 NA) was used. mVenus and tdTomato fluorescence were separated using a Chroma 565dcxr dichroic mirror and isolated using HQ510/70 (Chroma), and 630/92 (Semrock) or 635/90 (Chroma) emission filters, respectively.

For glutamate uncaging, 2.25 mM MNI-L-glutamate (Tocris Bioscience) was added to the ACSF. Glutamate uncaging was induced by applying 70 – 100 mW (measured at the back focal plane of the objective) light pulses of 0.2 – 0.5 ms duration 720 nm light. Approximately 60% of the power was transmitted though the objective. The power was held constant within the same field of view but was changed empirically depending on the tissue depth. The uncaging depth in the slice was restricted to 10 – 80 µm from tissue surface. To elicit uEPSCs the uncaging beam was positioned either at the tip of chosen spines or adjacent to the identified PSD-95^mVenus^ puncta. uEPSC amplitudes were measured by averaging 5 – 10 trials per position elicited at a frequency of 0.1 Hz. To reliably determine uEPSC amplitudes, traces were smoothened using a sliding 2ms window. Peak uEPSC amplitude was calculated as the difference between the peak current amplitude and mean current amplitude over a 25 ms window prior uncaging. Current amplitudes smaller than baseline + 3 times standard deviation were considered noise and were set to zero. To calculate charge transfer, a time window for integration was determined per dendrite. A mean onset time, time between 10% of the peak amplitude to peak, mean decay time, time between peak and 10 % of the peak amplitude, and mean peak time was calculate based on all uEPSCs recorded within a given dendrite. The integration window was placed between the mean onset time and mean decay time, relative to the mean peak time. The charge transfer was corrected using a baseline charge transfer, calculated using the same time integration window placed before uncaging.

#### Cellular Reconstructions

For reconstruction of cell morphology, we performed intracellular recording with biocytin in the recording pipet. The slices were overnight fixed in 4% paraformaldehyde and 4% sucrose in 0.1 M phosphate buffer (PB, pH 7.4). Slices were washed in PB for 60 minutes followed by a 40-minute treatment with 3% peroxide, to reduce endogenous peroxidase activity, and a 20-minute wash in PB. Biocytin-filled cells were visualized using an avidin-biotinylated horseradish peroxidase complex reaction (Vectastain-Elite, Vector Laboratories). Slices were incubated in 1% Vectastain-Elite with 0.5% triton in PB, first for 30 minutes at room temperature followed by overnight at 4 °C and 1.5 hours at room temperature. Slices were wash for 1 hour in PB and incubated in 1 mg/ml 3,3’-diaminobenzidine (DAB, Sigma-Aldrich) with 0.0002% CoCl_2_ and 0.0004% (NH_4_)_2_Ni(SO_4_)_2_ at 4 °C. Chromogenic reaction was started by adding peroxide (0.0003% end concentration) and kept incubating until cell morphology was clearly visible. The chromogenic reaction was stopped by extensive washing in PB. Slices were mounted on gelatinized microscope slides and left to dry overnight, or until dry, in an 80% humidity chamber. Subsequently, slices were dehydrated in steps of 10% EtOH from 10% to 90%, twice in 100% EtOH, and xylenes, each step for 10 minutes. Slices were embedded and coverslipped in Eukitt. Cellular morphology was reconstructated using Neurolucida (MBF bioscience) at 1000x magnification. Reconstructions were not corrected for shrinkage.

#### In-Utero Viral Transfection

For sparse labelling of layer 2/3 pyramidal neurons, we performed in-utero viral transfection as previously described (Fortin et al., 2014). Briefly, 1 µl of AAV2/1 virus expressing Cre under the Synapsin promoter was diluted to an empirically-determined level (1:20 – 1:50) optimal for sparse labeling of dendrites in layer 1. The diluted virus was injected into the lateral ventricles of E15.5 pregnant mice.

#### Cranial Window Surgery

Surgical implantation of the cranial window was performed similarly to Fortin et al., 2014, with several exceptions. Briefly, adult mice were anesthetized with isoflurane (induction: 4 percent in 1.0 l/min medical grade oxygen, maintenance: 1.5 percent in 0.2 L/min medical grade oxygen) and given Dexamethasone (20 µl of 4 mg/ml solution) and Buprenorphine (80 µl of 0.03 mg/ml solution) to mitigate post-surgical inflammation and pain, respectively. Using sterile procedures, the scalp was retracted, and a craniotomy was performed over the left barrel cortex (from bregma, 1.2 mm posterior, 3.4 mm lateral). The drilling site was irrigated with sterile cortex buffer (in mM: NaCl 125, KCl 5, Glucose 10, HEPES 10, CaCl_2_ 2, MgSO_4_ 2, pH 7.4) to clear bone dust from the site and to prevent overheating of the neuropil. After removing the flap of bone, a customized glass window (laser cut by Potomac Photonics, assembled in-house) was gently pushed into the craniotomy. The glass window consisted of three layers of glass fused with Norland Optical Adhesive 61. These layers consisted of the following: a 3.5 mm circular 1 coverslip on bottom, an inner ‘donut’ ring (ID: 3 mm, OD: 3.5 mm) in the middle, and an outer ‘donut’ ring (ID: 3mm, OD: 5 mm) on top. The ‘donut’ optical window assembly allows for high-resolution imaging through a single-piece of cover glass, and provides gentle pressure on the cerebral cortex to mitigate motion-induced artifacts during imaging. The glass window was sealed in place, along with a custom aluminum head-bar, using a mixture of cyanoacrylate glue and dental cement. The animal recovered from surgery in a heated cage and monitored until anesthesia had worn off. A second dose of Buprenorphine and Dexamethasone was administered 24 hours after the cranial-window surgery.

#### Intrinsic-Signal Imaging

Approximately one week after cranial window implantation, experimental mice were sedated with chlorprothixene (50 µl of 100 µg/ml solution) and lightly anesthetized with isoflurane (1 % in 0.2 l/min medical grade oxygen). Intrinsic signal imaging was performed as previously described (Ma et al., 2018; Masino et al., 1993), through the glass ‘donut’ apparatus. The subject’s whiskers were stimulated with a piezo actuator coupled to a large vinyl surface that could stimulate the majority of the subject’s whiskers. Controls were occasionally performed by performing the experiment without stimulating the piezo actuator; in these experiments, no intrinsic signal responses were ever observed. Confirmatory experiments (Supplemental Figure 1B) stimulated single barrels by loading a single whisker into a glass capillary tube coupled to the piezo.

#### Longitudinal in vivo Two-Photon Imaging

After allowing the animal to recover from the cranial window implantation surgery for at least two weeks, dendrites in layer I of barrel cortex (S1) were imaged using a custom-built two-photon microscope based on the MIMMS design (Janelia) for *in vivo* imaging. The time course of longitudinal two-photon imaging commenced at least 5-days after intrinsic signal imaging, to mitigate off-target effects of chlorprothixene and whisker stimulation. Throughout the experiment, the presence of barbering was monitored by imaging the whisker length of all animals; animals that were barbered during the course of the experiment were excluded from analysis. Animals were anesthetized with isoflurane and mounted under a custom-built two-photon microscope via an aluminum headbar installed during the craniotomy surgery. All imaging was performed through a Nikon 16x, 0.8 NA objective. On the first day, PSD-95^mVenus^+ and tdTomato+ dendrites were imaged in layer 1 in areas of the window that yielded barrel-responses during intrinsic signal imaging. Images of dendrites were acquired at 512 x 512 pixels, a scan speed of 2 ms/line, and a field-of-view of ∼45 micrometers. No averaging was performed during image acquisition, but we oversampled in the z dimension (0.8 µm/step) which allowed for running average in the z dimension. The laser power at specimen was held constant for all imaging-sessions of the same field of view and never exceeded 45 mW at sample.

### QUANTIFICATION AND STATISTICAL ANALYSIS

#### General Statement / Statistics

Data analyses were performed with custom programs written in MATLAB (Mathworks), Python 2.7 and 3.8 (https://www.python.org/downloads/release/python-380/), or R (https://cran.r-project.org). All programs are freely available (github.com/hzlab/synvivo). Sample sizes are indicated in text or figure legends and *: p ≤ 0.05, **: p ≤ 0.01, and ***: p ≤ 0.001). Unless indicated otherwise, statistical tests and distributional best-fits were computed via the scipy package in Python 3.6.

#### Four-Dimensional Synapse Scoring and ROI Generation

PSD-95mVenus puncta were identified manually and their persistence and dynamics were scored in custom MATLAB software that allowed for the simultaneous visualization of the same synapse in three dimensions over all 8 experimental time points (Supplemental Figure 2). Only dendrites with high imaging quality over all imaging sessions were kept for analysis. At each time point, PSD-95^mVenus^ puncta were classified as residing on the shaft of the dendrite or on a spine. To reduce noise associated with analyzing four-dimensional two-photon imaging datasets, we implemented a “two-day rule”: for a synapse to be scored as eliminated, it must be absent for two consecutive imaging sessions. Spiny synapses were defined as synapses that protruded laterally from the dendritic shaft, with a clear emanating structure evident from the tdTomato channel. Shaft synapses were identified as PSD-95^mVenus^ puncta that colocalized with the dendritic shaft. Due to the relatively poor axial resolution of two-photon microscopes, we only classified spines based on their geometry, relative to the dendrite, in the X-Y dimensions: all axially protruding spines were thus classified as shaft synapses.

After initial scoring for the presence and persistence of each synapse, ellipsoid ROIs were generated for each synapse and manually adjusted to ensure proper fit. Changes in the ROI size were mirrored across all imaging-days for a given synapse, ensuring that measures of changing intensity were not subject to changes in ROI size.

#### Weight Extraction

To compute the weight of a synapse, we used the elliptical ROIs generated manually in custom scoring software. Next, we searched in the axial-plane (within a defined window), to find a synapse’s center-of-mass and adjusted the z-position of the synaptic ROI if necessary. Next, the average pixel value of a user-designated background ROI was subtracted from each pixel within the ROI. Additionally, we subtracted tdTomato’s bleedthrough in the green collection channel (for mVenus). Experiments in tdTomato-only dendrites estimated this value at ∼2%, which was used for these experiments. Next, the background-and bleedthrough-subtracted pixels within an ROI were integrated across 5 axial-planes (±1.6 µm). These values were normalized to the average of the 40^th^ to 60^th^ percentile of all such intensities on a dendrite for a single day. This normalization assumed that the average synaptic strength along a dendritic segment is stable across dendrites and across days, but it was necessary to pool data from different dendrites and dates together due to different imaging conditions (depth, local blood vessels, etc), and day-to-day variations in imaging conditions (e.g., see Figure S3A and S3B). We used this approach over an alternative to normalize to the dendritic cytosolic marker fluorescence because the expression of PSD-95^mVenus^ and tdTomato might not be stoichiometric across cells, and mVenus and tdTomato were differentially bleached over days (Figure S3C and S3D).

#### Markov-Chain Model

##### Fitting Procedure

We binned the non-zero synaptic strengths into 40 equal-width bins, but group the last 21 bins into a single bin given that the counts were small in these last 20 bins. These bins constituted the “states” of the Markovian transition matrix, resulting in 21 total states (20 states of non-zero synaptic strength and one state reserved for a strength of 0). The number of bins were empirically determined. Other number of bins gave qualitatively similar results. The Markovian transition matrix was fit using a maximum likelihood estimator (MLE), in which for each pair of consecutive observation days (t and t+1), we counted the number of times the synaptic strength changed from state s_1_ on day t to state s_2_ on day t+1, for all pairs of states s_1_ and s_2_. We then normalized these counts to form a conditional probability distribution such that 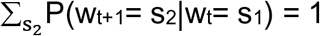. These transition probabilities were used as the elements of the Markov transition matrix. Each row of this matrix indicates the initial state s_1_ and each column indicates the final state s_2._ Thus each row is a conditional probability distribution of future state given current state and sums to 1.

##### Stationary Distribution

The stationary distribution was obtained from the right eigenvector of the Markov transition matrix with an eigenvalue of 1. This same stationary distribution can also be obtained from iterating the Markov dynamics many times from an arbitrary initial distribution. We compared this to the empirical stationary distribution fit using a maximum likelihood estimator (MLE), where the synaptic strengths were binned into the same states as used for the Markovian transition matrix, and the number of occurrences of synaptic strengths (aggregated across observation days) was counted in each bin, normalized by the total number of synapses.

To generate errorbars, the original data were sampled with replacement from 30 bootstrap runs. In each bootstrap run, the data were fit with a Markov transition matrix to obtain the stationary distribution from the eigenvector (model, as described above), as well as fit by the empirical stationary distribution (experimental).

##### Cross Correlation Coefficient of Weight Changes

Once the Markovian transition matrix had been established, we sampled from the chain starting at the stationary distribution. Although the Markovian transition matrix returned discrete states at each time point, we converted those states back to continuous synaptic strengths by uniformly sampling between the minimum and maximum synaptic strength value within that state bin. This procedure resulted in a continuous valued trajectory of synaptic strengths *S* for each synapse in the original dataset.

We then computed the cross-correlation coefficient on both *S* and the original dataset *W* (both of which were matrices of size, number of synapses N x total observation days T). First, the consecutive weight changes were computed in *W*, given by v_t_ = w_t_ − w_t-1_, resulting in the matrix V, and in *S*, v̂_t_ = s_t_ − s_t-1_, resulting in the matrix V̂. The cross correlation was then computed from these consecutive weight change matrices. For a given increment of observation time Δ,

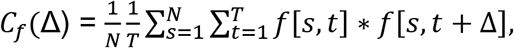

where *f* can either be the matrix V or the matrix V̂. The “cross-correlation coefficient” was given by *CC_f_*(Δ) = *C*_*f*_(Δ)/ *C*_*f*_(0), in order to attain a maximum value of 1.

##### Survival Fraction

For the simulated survival fraction in Figure 4K, we generated a continuous valued trajectory of synaptic strengths *S* from the Markovian transition matrix (as described above), but starting from the distribution of synaptic strengths from the first day to match the original data. The survival fraction was then computed both in the model and in the data as the fraction of synapses on each observation day that persisted from the original set of synapses on the first day. In the survival fraction simulations, synapses could only enter and exit the “0” state once to simulate synapse addition and elimination, respectively.

#### Kesten Model

A Kesten process is given by w_t+1_ = ε_t_·w_t_ + η_t_, which can be rewritten as w_t+1_ = w_t_ + (ε_t_ – 1)·w_t_ + η_t_, where w_t_ is the synaptic size at time *t* and ε_t_ and η_t_ are random variables that can be drawn from any distribution (we set them to be Gaussians, as explained below). Set α_t_ = ε_t_ – 1, so Δw_t_ = (ε_t_ – 1)·w_t_ + η_t_ = α_t_·w_t_ + η_t_. We assumed that α_t_ and η_t_ were random processes as they were the result of unobserved pre- and post-synaptic activity. We simply modeled them as Gaussians, whereby α_t_ ∼ N(a_t_, b_t_) and η_t_ ∼ N(c_t_, d_t_). Here N(a,b) denotes a Gaussian distribution with mean a and variance b. As a result, Δw_t_ ∼ N·(a_t_·w_t_ + c_t_, b_t_·w_t_^2^ + d_t_). We binned the weights w_t_ in order to get an approximate distribution for the synaptic strengths at day *t*, and then plotted the average value of the consecutive weight change <Δw_t_> = a_t_·w_t_ + c_t_, and its variance Var(Δw_t_) = b_t_·w_t_^2^ + d_t_, versus the average (across synapses in that bin), w_t_, and the square of this average, w_t_^2^, respectively. This allowed us to first determine respectively if a_t_, b_t_, c_t_, and d_t_ are indeed independent of momentary synaptic strength w_t_ (and can therefore be treated as constants in a linear regression), as well as their corresponding values.

To perform this linear regression, we binned the (nonzero) synaptic strengths (across synapses and observation days) into 20 bins with roughly equal numbers of elements, which for synapses onto the pyramidal cell type resulted in ∼165 synaptic strength values per bin, and for those onto the PV cell type resulted in ∼125 synaptic strength values per bin. We then computed Δw_t_ = w_t+1_ − w_t_, for all synaptic strength values w_t_ in that bin, from which we further computed <w_t_>, <w_t_>^2^, <Δw_t_>, and Var(Δw_t_) per bin. These four values per bin were then linearly regressed, namely <Δw_t_> vs. <w_t_> and Var(Δw_t_) vs. <w_t_>^2^ – yielding the coefficients a, b, c, and d, as described above.

---

**Figure S1. Related to Figure 1.**
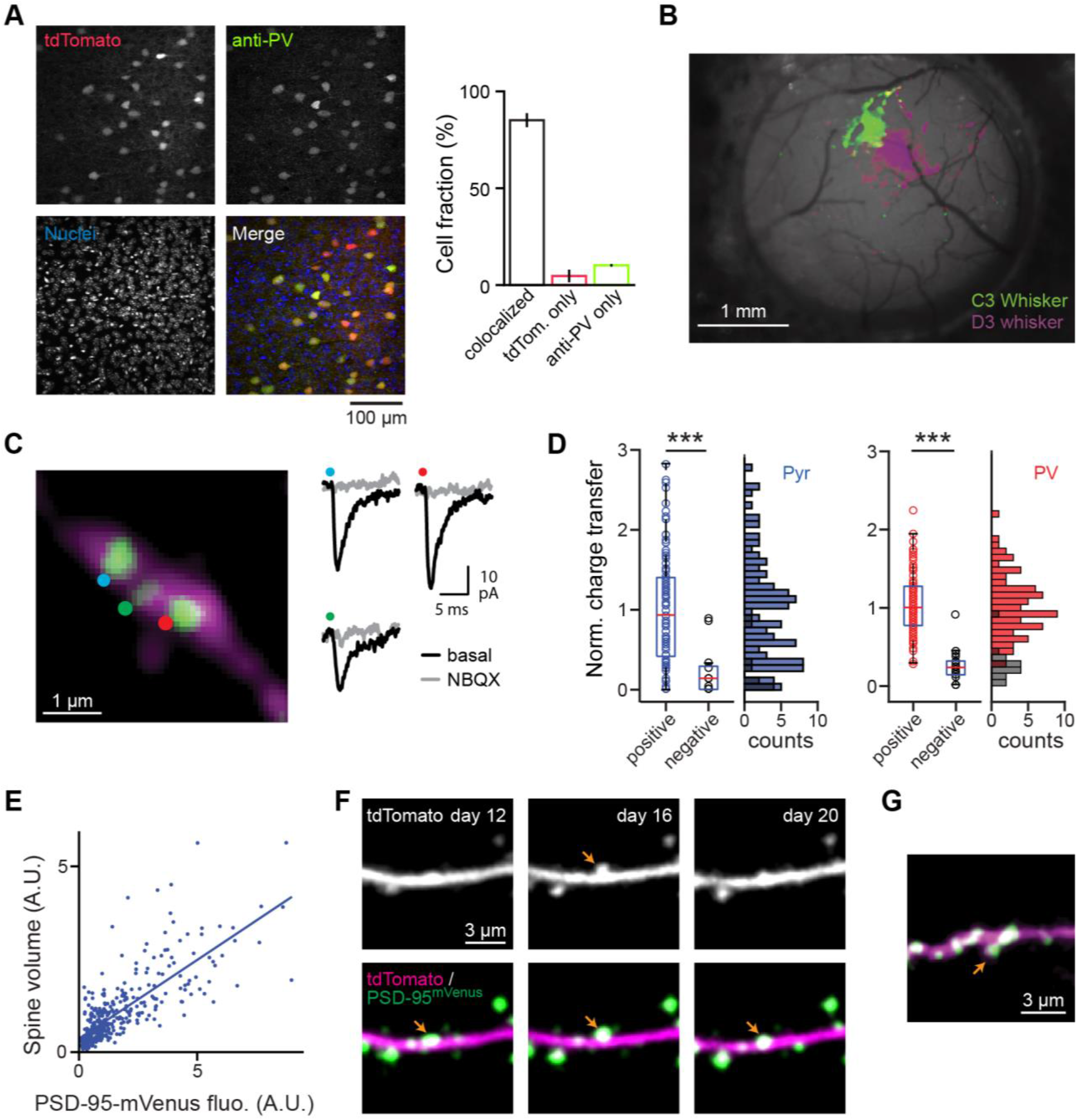
Validation and characterization of ENABLED-strategy for *in vivo* cell-type-specific targeting in the adult mouse barrel-cortex. (A) Immunohistochemical confirmation of the *PV-IRES-Cre* driver line in the barrel cortex of adult mice. Left: example image showing tdTomato fluorescence (expressed by the *Ai9* reporter line), anti-PV labelling (via immunostaining), and nuclei (Hoescht). Right: Percentage of all cells identified by the red (tdTomato) or green (anti-PV) fluorescence alone, or colocalized. Red and green images were analyzed independently. Out of all neurons identified in both channels, 85.1% were both tdTomato-positive and PV antibody-positive. Only 4.7% of tdTomato-positive neurons lacked immune labels for the PV protein. Considering the variability associated with antibody staining, this result indicates that the PV-Cre driver line reliably labels PV+ interneurons in the barrel cortex of adult mice. (B) Functional validation of barrel identity using intrinsic imaging in the barrel cortex. Two adjacent whiskers (C3 and D3, as indicated) were loaded into a capillary glass tube and stimulated sequentially while performing intrinsic-signal imaging. The resulting intrinsic-signal responses displayed a spatial shift that reflects the known preservation of mystacial organization in S1. (C) NBQX blocks uEPSCs. Representative image of a PV+ neuronal dendritic stretch with PSD-95^mVenus^ puncta. Two-photon glutamate uncaging elicited uEPSCs (black lines) corresponding to the uncaging locations (colored dots) in image. The uEPSC were blocked by the AMPA receptor antagonist NBQX (gray lines). _(D)_ Charge transfer of uEPSC responses normalized to the averaged response of each stretch of dendrites, and their histograms, at locations with (colored circles and bars) and without (black circles and bars) PSD-95^mVenus^ puncta (see Figures 1D and 1E for examples). Glutamate uncaging at PSD-95-positive puncta yielded much higher uEPSC than locations without PSD-95 in both Pyr and PV neurons (Mann-Whitney test: for Pyr: n_w/ PSD-95_ = 101 and n_w/o PSD-95_ = 10 uncaging spots, *U* = 5612, *P* < 0.001; for PV: n_w/ PSD-95_ = 75 and n_w/o PSD-95_ = 16 uncaging spots, *U* = 3241, *P* < 0.001). Tukey-style boxplots represent the median (red line), first and third quartiles (box), and lowest/highest data within 1.5 times interquartile range (whiskers). n (dendrites / cells / mice) = 9 / 6 / 3 for pyramidal neurons and 11 / 9 / 4 for PV+ neurons. (E) Correlation of spine volume with PSD-95^mVenus^ fluorescence intensity *in vivo* for spines clearly separated from their parental dendritic shafts (n = 361 spines from 14 L2/3 pyramidal dendrites). (F) PSD-95^mVenus^ permits the longitudinal visualization of axially-protruding spines. Representative images from three consecutive image sessions of a stretch of L23 pyramidal dendrite. Both morphology-only (top) and morphology with PSD-95^mVenus^ (bottom) are shown. In the morphology-only channel, a representative spine (orange arrow) can only be visualized for a single time-point; however, inclusion of the PSD-95^mVenus^ channel reveals its persistence throughout all 3 imaging sessions. This is likely the result of the spine rotating into the axial-plane and colocalizing with the dendritic shaft. Such phenomena can lead to systematic over-estimation of synaptic dynamics when using spine-morphology as the sole proxy for the presence of a synapse. (G) Representative image of a spine (arrow) on a PV+ dendrite. Such structures were infrequently observed *in vivo*.

**Figure S2. Related to Figure 3.**
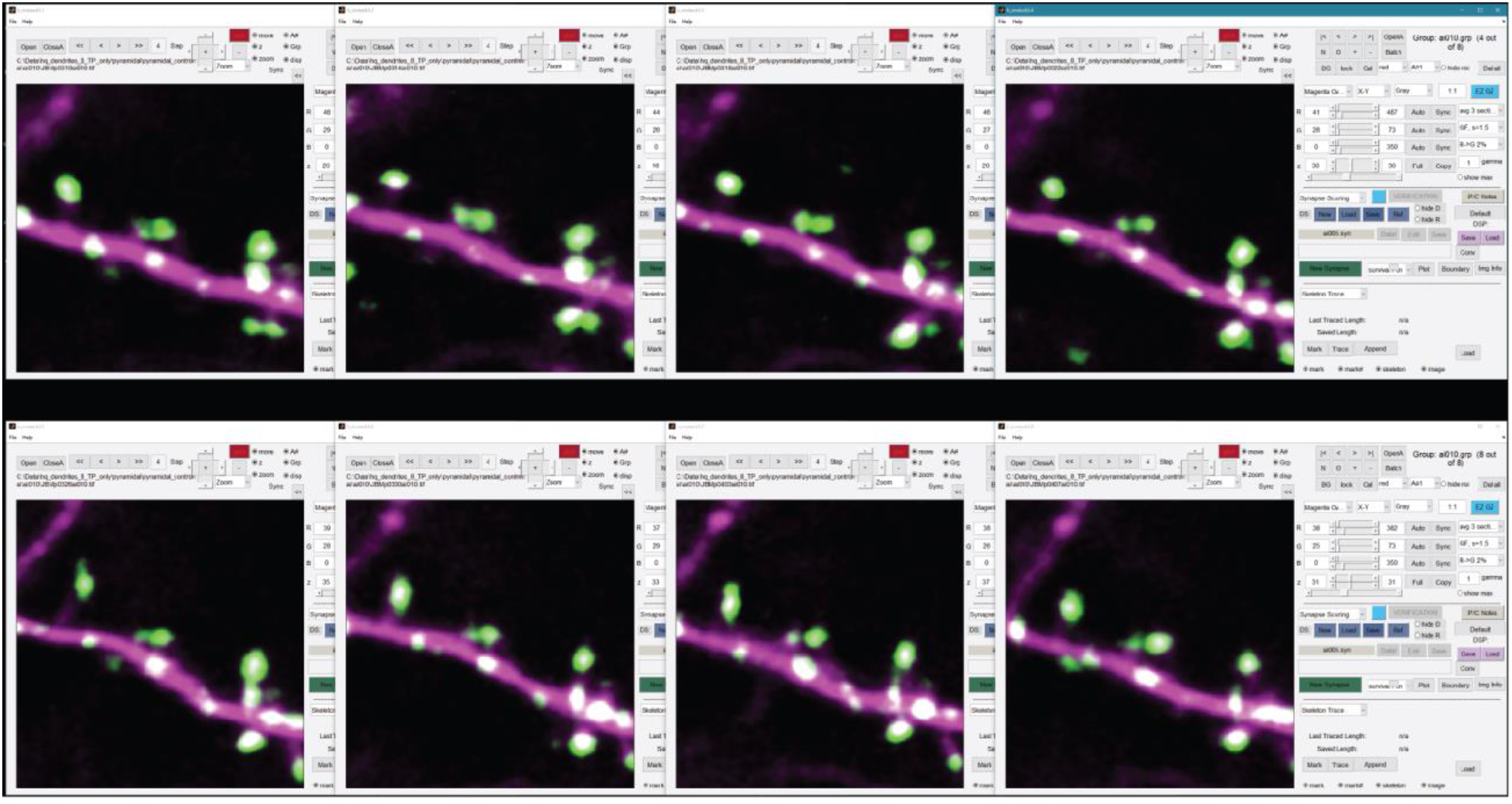
Representative screenshot of custom software for scoring multi-day longitudinal data side-by-side. Longitudinal *in* vivo two-photon data consist of an arbitrary number of three-dimensional images of the same field-of-view. For datasets with more than two time points, (as in this study), iterative pairwise registration of synaptic structures is tedious and can result in accumulated errors when going through different time points. In order to minimize such errors, we developed a custom scoring GUI in MATLAB that permits the simultaneous visualization of the same field-of-view from an arbitrary number of experimental time points (hardware permitting). Using intuitive keyboard hotkeys and mouse-inputs, the four-dimensional dataset can be explored simultaneously with ease by, for example, navigating in X-Y with arrow keys (for all time points) and changing the axial plane with the scroll wheel. To minimize scoring errors, synapses were scored one-at-a-time starting with first appearance, with a mechanism to bring the user’s cursor to the same X-Y coordinate at each subsequent time point (as an additional means of determining synaptic identity). Puncta could be classified as protruding on a spine or colocalizing with the dendritic shaft, or flagged to indicate that the scorer was unsure about the identity of a given synapse on a given day.

**Figure S3. Related to Figure 3.**
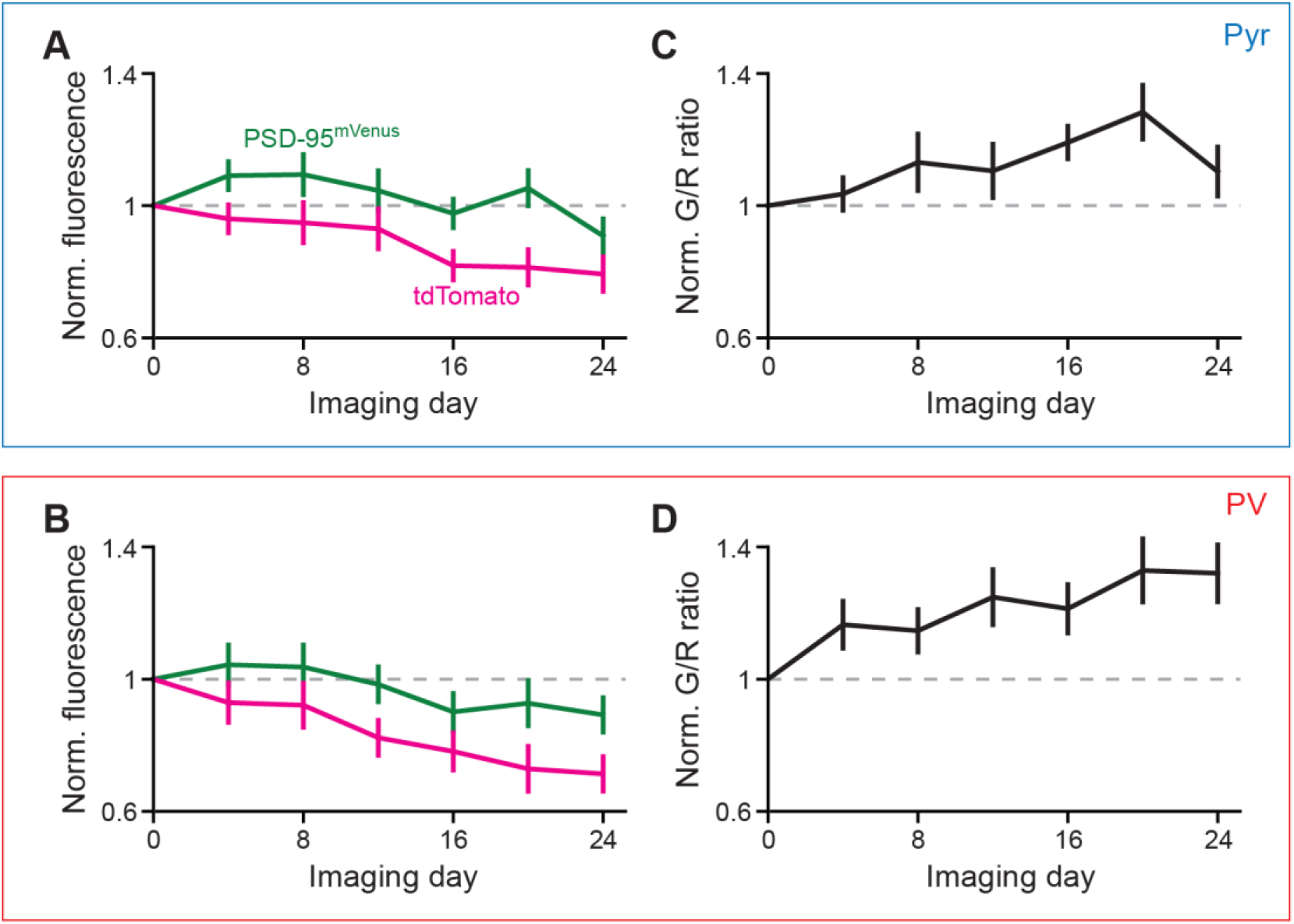
TdTomato and PSD-95^mVenus^ proteins are differentially bleached. (A, B) Time course of the average raw PSD-95^mVenus^ and tdTomato fluorescence values on L23 pyramidal dendrites (A) and PV+ dendrites (B) *in vivo*, normalized to the average raw-values on the first day of imaging. (C, D) Time course of the average (40^th^ to 60^th^ percentile) green-to-red ratio of puncta on L23 pyramidal dendrites (C) and PV+ dendrites (D) *in vivo*, normalized to the average green-to-red ratio on the first day of imaging. Note: the accumulated bleaching of cytosolic tdTomato and PSD-95^mVenus^ is not equivalent: tdTomato fluorescence attenuates more rapidly and is more attenuated by the 7^th^ imaging time point. These data suggest that the green-to-red ratio of a given synapse, as a normalized quantification of green intensity, is prone to artifact.

**Figure S4. Related to Figure 4.**
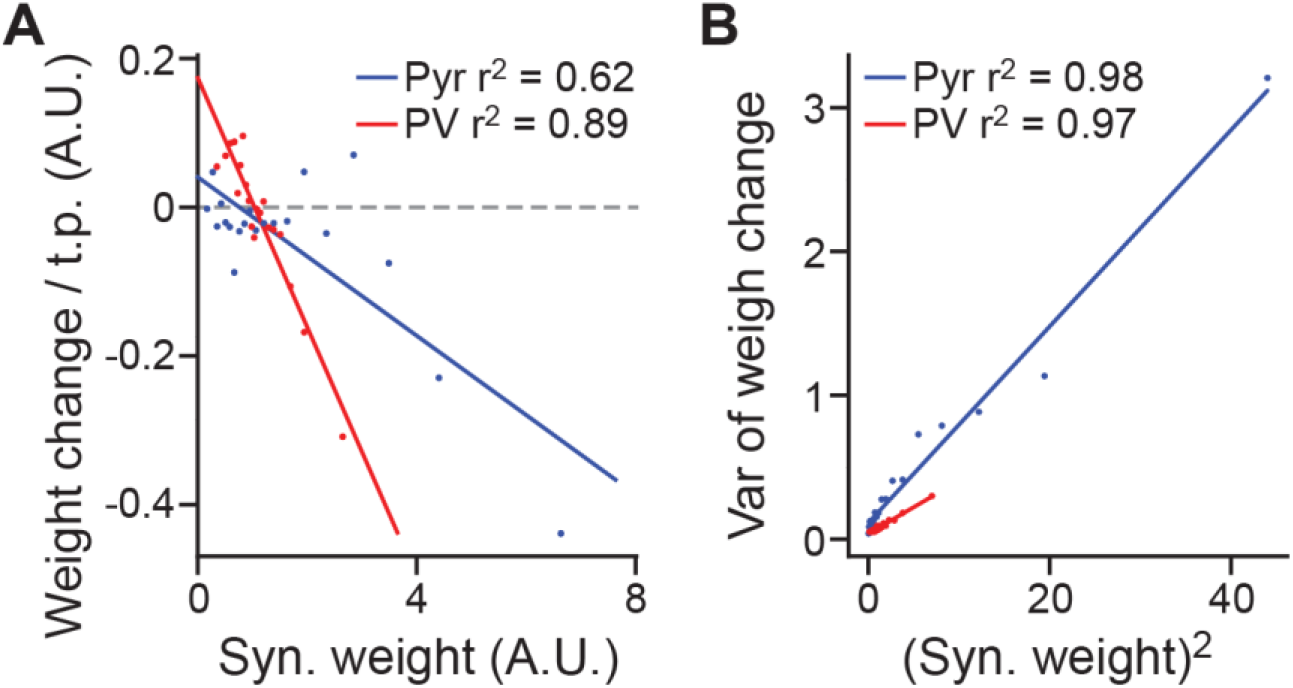
Synaptic weight dynamics are cell type-specific. (A) For each cell type, synaptic strengths were binned into 20 bins with equal counts. A line to the average (within bins) consecutive weight changes w_t+1_ – w_t_ was fit using the average synaptic weight <w_t_> as input (See **STAR METHODS**\***Kesten Model*** for details). Following the framework of Ziv and Brenner, 2018 that describes weight-dynamics as a combination of addition and weight-dependent multiplicative changes, these data suggest that excitatory synapses on PV+ dendrites have a stronger additive component in their dynamics than those on L23 pyramidal dendrites. (B) Same as A, but the variance of the consecutive weight changes w_t+1_ – w_t_ was fit using the square of the average synaptic weight <w_t_>^2^ as input.

